# Axon morphology and intrinsic cellular properties determine repetitive transcranial magnetic stimulation threshold for plasticity

**DOI:** 10.1101/2023.09.25.559399

**Authors:** Christos Galanis, Lena Neuhaus, Nicholas Hananeia, Zsolt Turi, Peter Jedlicka, Andreas Vlachos

**Author notes:** Correspondence to, Department of Neuroanatomy, Institute of Anatomy and Cell Biology, Faculty of Medicine, University of Freiburg, Freiburg 79104,Germany.

## Abstract

Repetitive transcranial magnetic stimulation (rTMS) is a widely used therapeutic tool in neurology and psychiatry, but its cellular and molecular mechanisms are not fully understood. Standardizing stimulus parameters, specifically electric field strength and direction, is crucial in experimental and clinical settings. It enables meaningful comparisons across studies and facilitating the translation of findings into clinical practice. However, the impact of biophysical properties inherent to the stimulated neurons and networks on the outcome of rTMS protocols remains not well understood. Consequently, achieving standardization of biological effects across different brain regions and subjects poses a significant challenge. This study compared the effects of 10 Hz repetitive magnetic stimulation (rMS) in entorhino-hippocampal tissue cultures from mice and rats, providing insights into the impact of the same stimulation protocol on similar neuronal networks under standardized conditions. We observed the previously described plastic changes in excitatory and inhibitory synaptic strength of CA1 pyramidal neurons in both mouse and rat tissue cultures, but a higher stimulation intensity was required for the induction of rMS-induced synaptic plasticity in rat tissue cultures. Through systematic comparison of neuronal structural and functional properties and computational modeling, we found that morphological parameters of CA1 pyramidal neurons alone are insufficient to explain the observed differences between the groups. However, axon morphologies of individual cells played a significant role in determining activation thresholds. Notably, differences in intrinsic cellular properties were sufficient to account for the 10 % higher intensity required for the induction of synaptic plasticity in the rat tissue cultures. These findings demonstrate the critical importance of axon morphology and intrinsic cellular properties in predicting the plasticity effects of rTMS, carrying valuable implications for the development of computer models aimed at predicting and standardizing the biological effects of rTMS.

## INTRODUCTION

Repetitive transcranial magnetic stimulation (rTMS) is a non-invasive technique that modulates cortical excitability beyond the stimulation period [1–3]. Despite its increasing use for treating neuropsychiatric disorders such as major depression [4–8], the cellular and molecular mechanisms of rTMS in human cortical networks remain not well understood [9,10]. Animal models, both *in vivo* and *in vitro,* have provided important insights into mechanisms by which rTMS modifies neuronal circuit excitability and plasticity [11–16]. It has been shown for example that rTMS affects the functional and structural properties of excitatory and inhibitory synapses [11,13,17], and that it facilitates the reorganisation of abnormal cortical circuits [18,19]. Recently, experimental evidence for an involvement of microglia, the brains resident immune cells in rTMS-induced synaptic plasticity was provided [16].

Although rTMS has shown robust neurobiological effects in animal models, its efficacy in humans varies significantly [20–23] due to challenges in dose standardization, among others [24,25]. Considerable effort has been made to standardize the electric field strength across brain regions and subjects to improve reproducibility and better understand the effects of single pulse and rTMS across brain regions [26–28]. Meanwhile, it is becoming increasingly clear that computational models that predict the strength and orientation of TMS-induced electric field must be extended to biological effects, i.e., the electric fields must be coupled to biophysically realistic models [29,30]. Indeed, these computational approaches provided important insight into the role of neuronal morphologies, specifically axons and myelination, which seem to play a critical role for single pulse TMS [31]. Although some attempts have been made to compute rTMS-induced changes in intracellular calcium levels, which could be used to predict plasticity effects [30], our knowledge regarding dose-response interrelation of rTMS-induced synaptic plasticity is limited. As a consequence, it is currently also not possible to compute and standardize synaptic plasticity induction across brain regions and subjects.

In this study, we explored the effects of 10 Hz repetitive magnetic stimulation (rMS) on CA1 pyramidal neurons of mouse and rat entorhino-hippocampal slice cultures. We discovered that rat CA1 pyramidal neurons required a 10% stronger intensity (MSO, maximum stimulator output) than mice to express plasticity. Our results suggest that axon morphologies and intrinsic cellular properties are key determinants of rTMS-induced plasticity. These insights carry important implications for the standardization of the “biological dose” of rTMS in comparable neuronal networks, both within and across subjects.

## MATERIALS AND METHODS

### Ethics statement

Mice and rats were maintained in a 12-hour light/dark cycle with food and water ad libitum. Every effort to minimize the distress and pain of animals was made. All experimental procedures were performed according to the German animal welfare legislation, approved by the appropriate animal welfare committee and the animal welfare officer of the University of Freiburg.

### Animals

Mice of the strain C57BL6/J and rats of the strain Wistar (Crl:WI) of both sexes were used in this study.

### Preparation of organotypic tissue cultures

Organotypic tissue cultures containing the hippocampus and the entorhinal cortex were prepared at postnatal day 3-5 from mice and rats of either sex as described preciously [11,32].

### rMS in vitro

Tissue cultures were transferred in a standard 35 mm petri dish filled with standard extracellular solution (129 mM NaCl, 4 mM KCl, 1 mM MgCl2, 2 mM CaCl2, 4.2 mM glucose, 10 mM HEPES, 0.1 mg/ml streptomycin, 100 U/ml penicillin, pH 7.4; preheated to 35 °C; 365mOsm with sucrose). A 70-mm figure-of-eight coil (D70 Air Film Coil, Magstim) connected to a Magstim Super Rapid2 Plus1 (Magstim) was placed 1 mm above the lid of the petri dish and the cultures were stimulated with a protocol consisting of 900 pulses at 10 Hz. Tissue cultures were orientated in a way that the induced electric field within the tissue was approximately parallel to the dendritic tree of CA1 pyramidal neurons. Species- and time-matched cultures were not stimulated, but otherwise identically treated served as the controls.

### Whole-cell voltage-clamp recordings

Whole-cell voltage-clamp recordings of CA1 pyramidal cells were conducted as previously described [11,13,32]. Neurons were recorded at a holding potential of −70 mV. Series resistance was monitored in 2 – 4 min intervals and recordings were discarded if the series resistance reached ≥30 MΩ and the leak current changed significantly.

### Whole-cell current-clamp recordings

Whole-cell current-clamp recordings of CA1 pyramidal cells were conducted at 35°C. The bath solution contained 126 mM NaCl, 2.5 mM KCl, 26 mM NaHCO3, 1.25 mM NaH2PO4, 2 mM CaCl2, 2 mM MgCl2, 10 mM glucose, 10 μM D-APV, 10 μM CNQX, and 10 μM bicuculline methiodide and was saturated with 95% O2/5% CO2. Patch pipettes contained 126 mM K-gluconate, 4 mM KCl, 4 mM ATP-Mg, 0.3 mM GTP-Na2, 10 mM PO-creatine, 10 mM HEPES, and 0.1% (w/v) biocytin (pH 7.25 with KOH, 290 mOsm with sucrose). Neurons were hyperpolarized with −100 pA and then depolarized up to +400 pA with 1- s-long 10 pA current injection steps. Recordings were discarded of the series resistance reached ≥15 MΩ.

### High-density microelectrode array (HD-MEA) recordings

HD-MEA recordings of mouse and rat tissue cultures were conducted at 35°C. The bath solution was similar to the one used for voltage-clamp recordings without the addition of any drugs. Cultures were placed on an Accura HD-MEA chip (3Brain, Switzerland) and acclimatized for 2 min before recording. Each tissue culture was recorded for 10 min with a BioCAM DupleX (3Brain, Switzerland).

### Neuronal filling, post hoc staining and imaging

CA1 pyramidal neurons were patched with pipettes containing 126 mM K-gluconate, 4 mM KCl, 4 mM ATP-Mg, 0.3 mM GTP-Na2, 10 mM PO-creatine, 10 mM HEPES, and 1% (w/v) biocytin (pH 7.25 with KOH, 290 mOsm with sucrose). The neurons were kept in the whole-cell configuration for at least 10 min during which they were depolarized with 100 ms current injections of 200 pA at 5 Hz.. Tissue cultures were fixed in a solution of 4% (w/v) PFA and 4% (w/v) sucrose in 0.01 M PBS for 1 h and further processed and images as previously described [32].

### Neuronal reconstructions

CA1 pyramidal cells were reconstructed using Neurolucida 360 (ver. 2019.1.3; MBF Bioscience) as described previously [30].

### Electric field modeling

Finite element method was used to create a three-dimensional mesh model consisting of two compartments, representing the bath solution and tissue cultures. The physical dimensions of the mesh model were based on the physical parameters of the *in vitro* settings, with a coil-to-Petri dish distance of 1 mm and the coil positioned above the culture. Electrical conductivities of 1.654 S/m and 0.275 S/m were assigned to the bath solution and culture respectively. The rate of change of the coil current was set to 1.4 A/ms at 1% MSO and scaled up to higher stimulation intensities. Simulations of macroscopic electric fields were performed using SimNIBS (3.2.6) and MATLAB (2023a). A validated 70 mm MagStim figure-of-eight coil was utilized in all simulations [33]. The 99th percentile of the E-field, which represents the robust maximum value, was extracted from the volume compartment of the tissue culture.

### Single cell modelling

Reconstructions were imported into the NeMo-TMS pipeline and endowed with a Jarsky model [34]. When axons are “swapped”, the original axon is removed from the cell at the point of intersection with the soma or dendrite, and replaced with the axon of another cell that has been severed at the same point. Each cell is oriented with the apical dendrite pointing in the positive y direction, and axon orientations relative to this are preserved in the swapping process. For single-cell simulations, TMS is simulated as a uniform electric field of varying intensity, with the threshold defined as the smallest TMS amplitude that elicits a somatic action potential.

### Experimental design and statistical analysis

Analyses were performed with the person analyzing the data blind to the experimental condition. For this project, we used one or two tissue cultures from each animal. Electrophysiological data were analyzed using pClamp 11.2 software suite (Molecular Devices), the Easy Electrophysiology 2.5.0.2 (Easy Electrophysiology Ltd.) and BrainWave (3Brain) software. Statistical comparisons were made using Mann–Whitney test (to compare two groups) two-way ANOVA and Kruskal-Wallis test as indicated in the figure captions and text (GraphPad Prism 7). p values of <0.05 were considered a significant difference. All values represent mean ± SEM.

### Digital illustrations

Confocal image stacks were exported as 2D projections and stored as TIFF files. Figures were prepared using Photoshop graphics software (Adobe). Image brightness and contrast were adjusted.

## RESULTS

### 10 Hz repetitive magnetic stimulation induces plasticity of excitatory and inhibitory synapses in mouse CA1 pyramidal neurons

A 10 Hz stimulation protocol consisting of 900 pulses at 50% MSO was used to assess the effects of rMS on synaptic plasticity in brain tissue cultures prepared from mice of either sex (Fig. 1A-C). Individual CA1 pyramidal neurons were patched and AMPA receptor mediated mEPSCs were recorded 2 – 4 h after stimulation. In line with our previous work [c.f., [11,16,35,36]] a significant increase in mean mEPSC amplitude was observed as compared to age-/time-matched control cultures that were treated in the exact same way except for rMS (control; Figure 1D-E).

**Figure 1:**
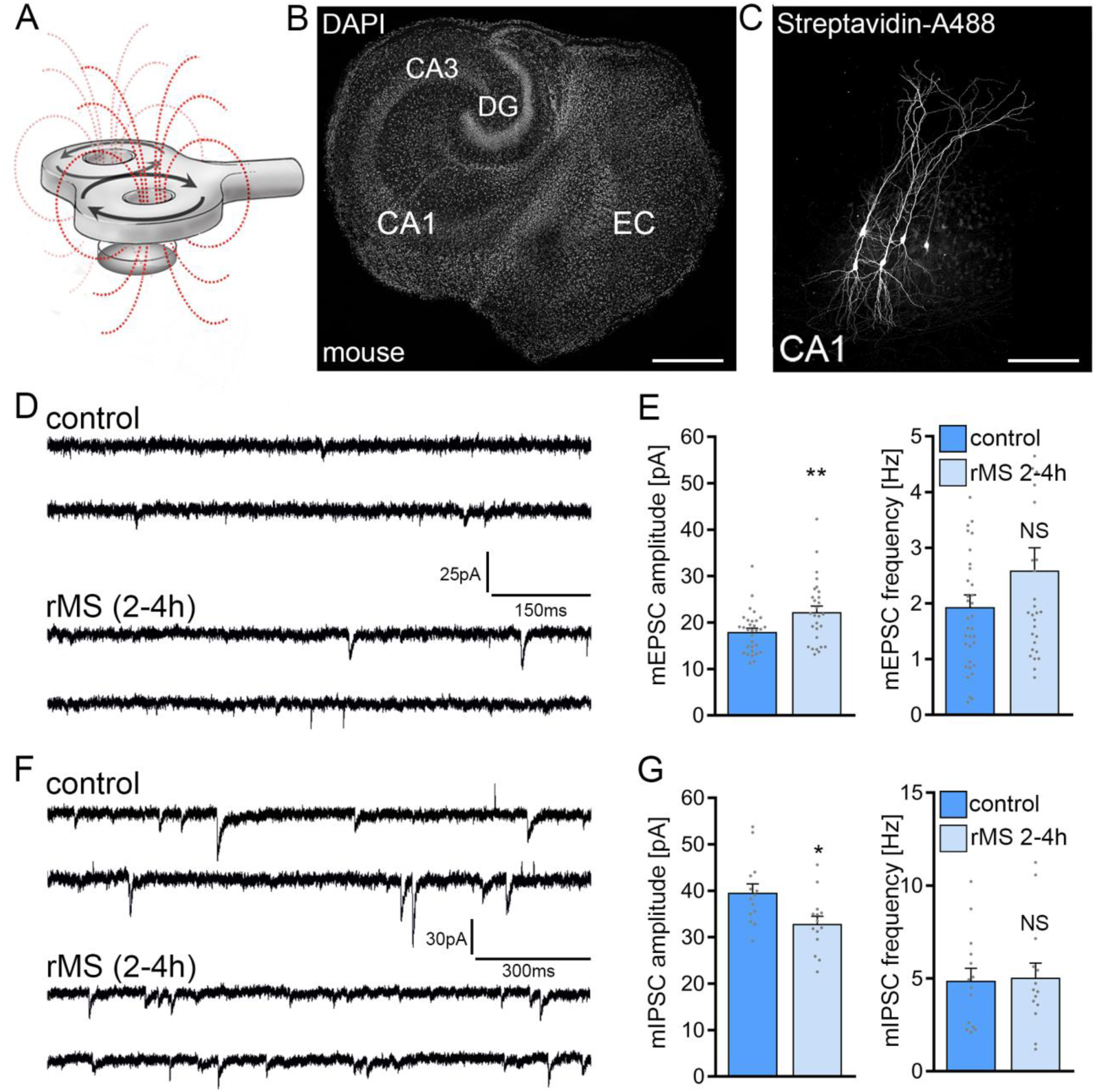
10 Hz repetitive magnetic stimulation (rMS) at 50% MSO induces synaptic plasticity in mouse CA1 pyramidal neurons. **(A)** Schematic illustration of the experimental setting. Organotypic tissue cultures are stimulated in a standard 35 mm petri dish filled with extracellular solution using a 70 mm figure-of-eight coil (900 pulses, 10 Hz, at 50 % maximum stimulator output). **(B)** Overview of an organotypic tissue culture. DAPI nuclear staining was used for visualization of cytoarchitecture. DG, Dentate gyrus; EC, entorhinal cortex; CA1 and CA3, *Cornu Ammonis* areas 1 and 3. Scale bar, 500 μm. **(C)** Patched CA1 pyramidal neurons filled with biocytin and identified *post hoc* with streptavidin-A488. Scale bar, 50 μm. **(D, E)** Sample traces and group data of AMPA receptor mediated miniature excitatory post synaptic currents (mEPSCs) recorded from mouse CA1 pyramidal neurons in sham-(control) and rMS-stimulated cultures 2 – 4 h after stimulation (control, n = 31 cells; rMS, n = 28 cells; Mann–Whitney test). **(F, G)** Sample traces and group data of GABA receptor mediated miniature inhibitory post synaptic currents (mIPSCs) recorded from mouse CA1 pyramidal neurons in sham-(control) and rMS-stimulated cultures 2 – 4 h after stimulation (control, n = 14 cells; rMS, n = 14 cells; Mann–Whitney test). Individual data points are indicated in this and the following figures by gray dots. Data are mean ± SEM. NS, Not significant. *p < 0.05. **p < 0.01.

In a different set of cultures, we assessed 10 Hz rMS-induced changes in GABA receptor mediated mIPSCs onto CA1 pyramidal neurons using the experimental approach described above. A reduction in mean mIPSC amplitude was observed in these experiments as reported in our previous study [(Fig. 1F-G); c.f., [13]]. These results confirm the robust effects of 10 Hz rMS on mEPSC and mIPSC amplitudes of CA1 pyramidal neurons of entorhino-hippocampal tissue cultures, which are consistent with a potentiation of excitatory synapses and a depression of inhibitory synapses.

### 10 Hz repetitive magnetic stimulation at 50% MSO does not affect synaptic strength in rat CA1 pyramidal neurons

The effects of the same 10 Hz protocol (10 Hz, 900 pulses, 50% MSO) were tested in tissue cultures prepared from rat brains (Fig. 2). Age-matched rat tissue cultures displayed a larger cross-section than mouse tissue cultures (Fig. 2A), without any apparent morphological differences in CA1 pyramidal neurons (Fig. 2B). Recordings of AMPA receptor mediated mEPSCs from CA1 pyramidal neurons showed no statically significant differences between control and 10 Hz rMS-stimulated preparations (Fig. 2A-B). Inhibitory synaptic strength was also unaffected, as no significant differences in mean mIPSC amplitude and frequency were detected 2 – 4 h after stimulation (Fig. 2C-D).

**Figure 2:**
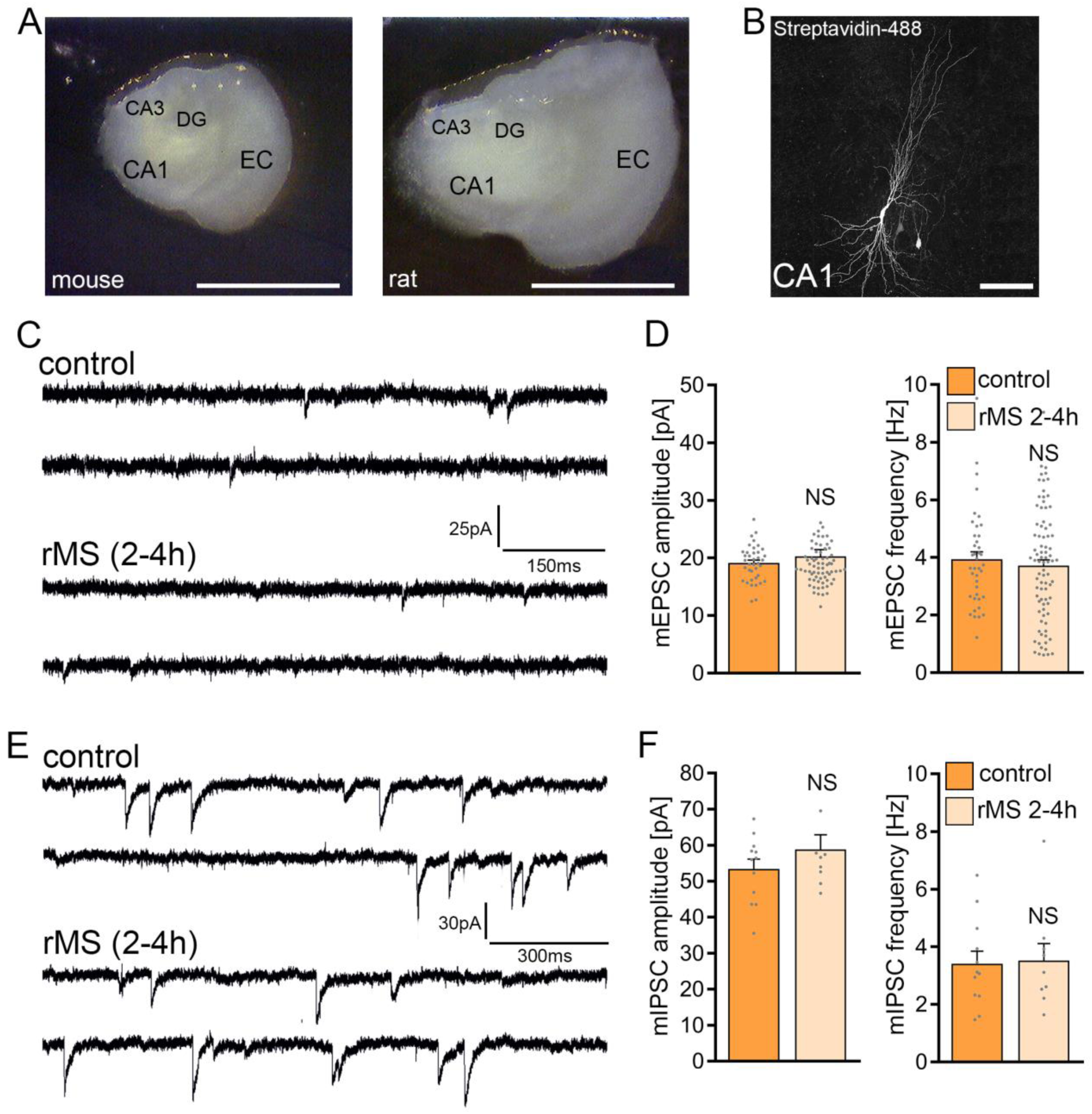
10 Hz repetitive magnetic stimulation (rMS) at 50 % maximum stimulator output does not affect synaptic transmission in rat CA1 pyramidal neurons. **(A)** Overview images of a mouse and rat organotypic tissue culture. DG, Dentate gyrus; EC, entorhinal cortex; CA1 and CA3, *Cornu Ammonis* areas 1 and 3. Scale bar, 500 μm. **(C)** Patched rat CA1 pyramidal neurons filled with biocytin and identified *post hoc* with streptavidin-A488. Scale bar, 50 μm. **(C, D)** Sample traces and group data of AMPA receptor mediated mEPSCs recorded from rat CA1 pyramidal neurons in sham-(control) and rMS-stimulated cultures 2 – 4 h after stimulation (control, n = 38 cells; rMS, n = 71 cells; Mann–Whitney test). **(E, F)** Sample traces and group data of GABA receptor mediated miniature inhibitory post synaptic currents (mIPSCs) recorded from rat CA1 pyramidal neurons in sham-(control) and rMS-stimulated cultures 2 – 4 h after stimulation (control, n = 12 cells; rMS, n = 9 cells; Mann–Whitney test). Data are mean ± SEM. NS, Not significant.

### Macroscopic electric field simulations reveal distinct maximum electric fields generated in mouse and rat tissue cultures

The electric field (E-field) strength induced in the mouse and rat slice cultures was described using computational modeling. Three-dimensional mesh models were created with two compartments (i.e., bath solution and slice cultures) using the finite element method (Fig. 3A). The physical dimensions of the mesh models were adapted from data obtain in mouse and rat brain issue cultures (Fig. 3B). Macroscopic modeling of the E-field revealed that stimulation at 50% MSO induces a stronger electric field in the mouse (20.4 V/m) when compared to the rat tissue culture (19.3 V/m). Based on the modeling we determined that 53 % MSO stimulation of rat tissue cultures would result in an E-field that is comparable to what we estimated in the mouse tissue cultures stimulated with 50 % MSO (Fig. 3C). Accordingly, another set of rat tissue cultures was stimulated with 53% MSO (10 Hz, 900 pulses) and AMPA-receptor mediated mEPSCs were recorded from CA1 pyramidal neurons 2 – 4 h after stimulation. No significant differences in mean mEPSC amplitude and frequency were observed in these experiments (Fig. 3D). We conclude that simulation-based standardization of electric fields may not suffice to achieve comparable biological effects in mouse and rat CA1 pyramidal neurons, i.e., in neurons embedded in networks with comparable architectures and properties.

**Figure 3:**
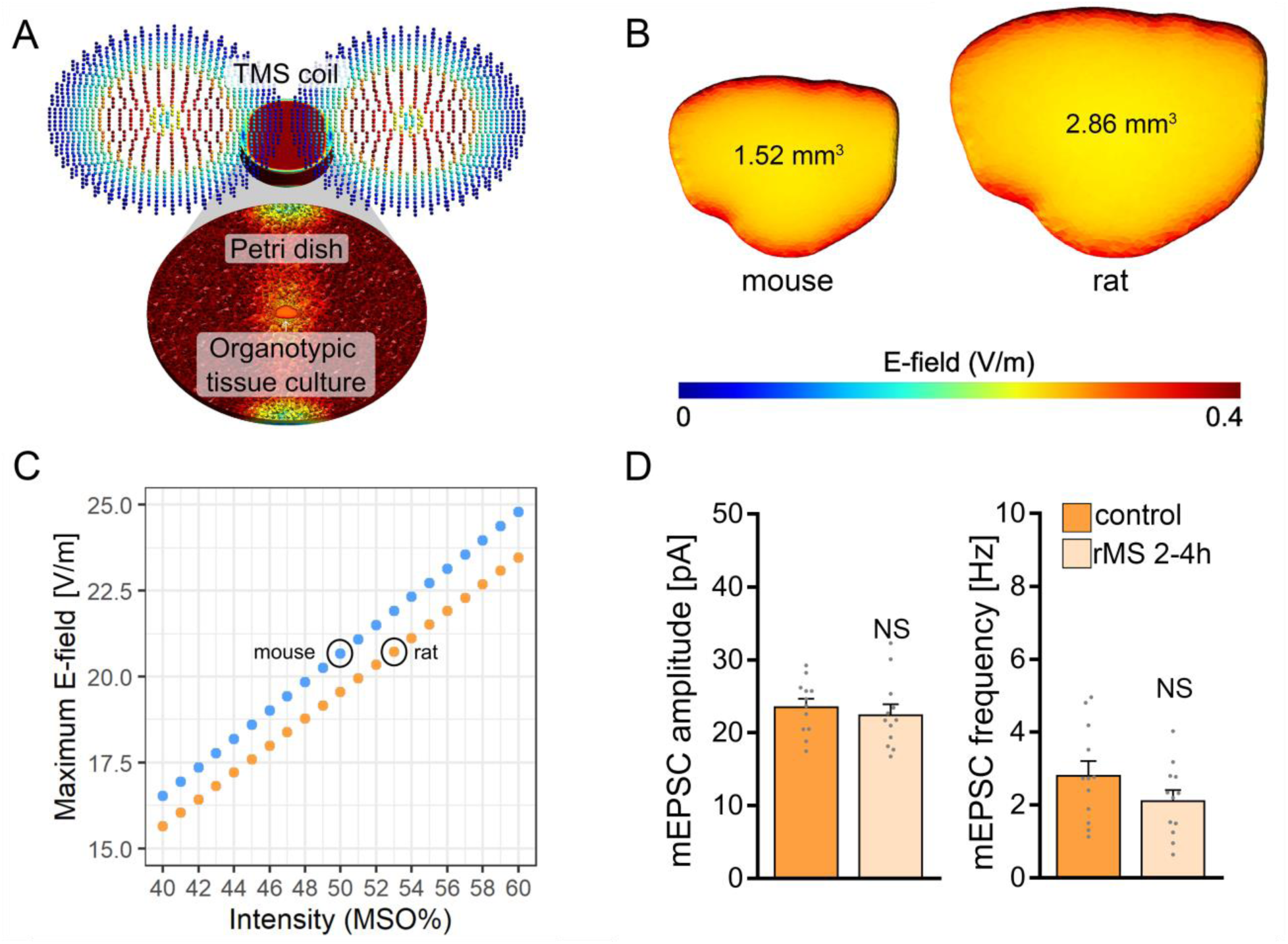
Modeling of repetitive magnetic stimulation (rMS)-induced electric fields in mouse and rat tissue cultures. **(A)** Visualization of the macroscopic electric field simulations generated by rMS *in vitro*. **(B)** Three-dimensional mesh models of mouse and rat organotypic tissue cultures and the electric fields generated by a single-pulse of rMS, respectively. **(C)** Comparison of the maximum electric field generated at distinct stimulation intensities in mouse and rat tissue cultures. The electric field generated in mouse slice cultures at 50 % maximum stimulator output is attained with 53% maximum stimulator output in rat tissue cultures. **(D)** Group data of AMPA receptor mediated mEPSCs recorded from rat CA1 pyramidal neurons in sham-(control) and rMS-stimulated cultures stimulated with 53 % maximum stimulator output and recorded 2 – 4 h after stimulation (control, n = 12 cells; rMS, n = 12 cells; Mann–Whitney test). Data are mean ± SEM. NS, Not significant.

### Baseline network activity is not significantly different between mouse and rat tissue cultures

To test for differences in spontaneous network activity between mouse and rat entorhino-hippocampal slice cultures basal firing rates and field potential rates were recorded in a different set of 3-week-old mouse and rat tissue cultures using high-density microelectrode array recordings (Fig. 4A, B). No significant differences between mouse and rat tissue cultures were observed in firing and field potential (FP) rates in these experiments (Fig. 4C-F). We conclude that baseline network activity is not responsible for the inability of rMS to induce plasticity in rat CA1 pyramidal neurons.

**Figure 4:**
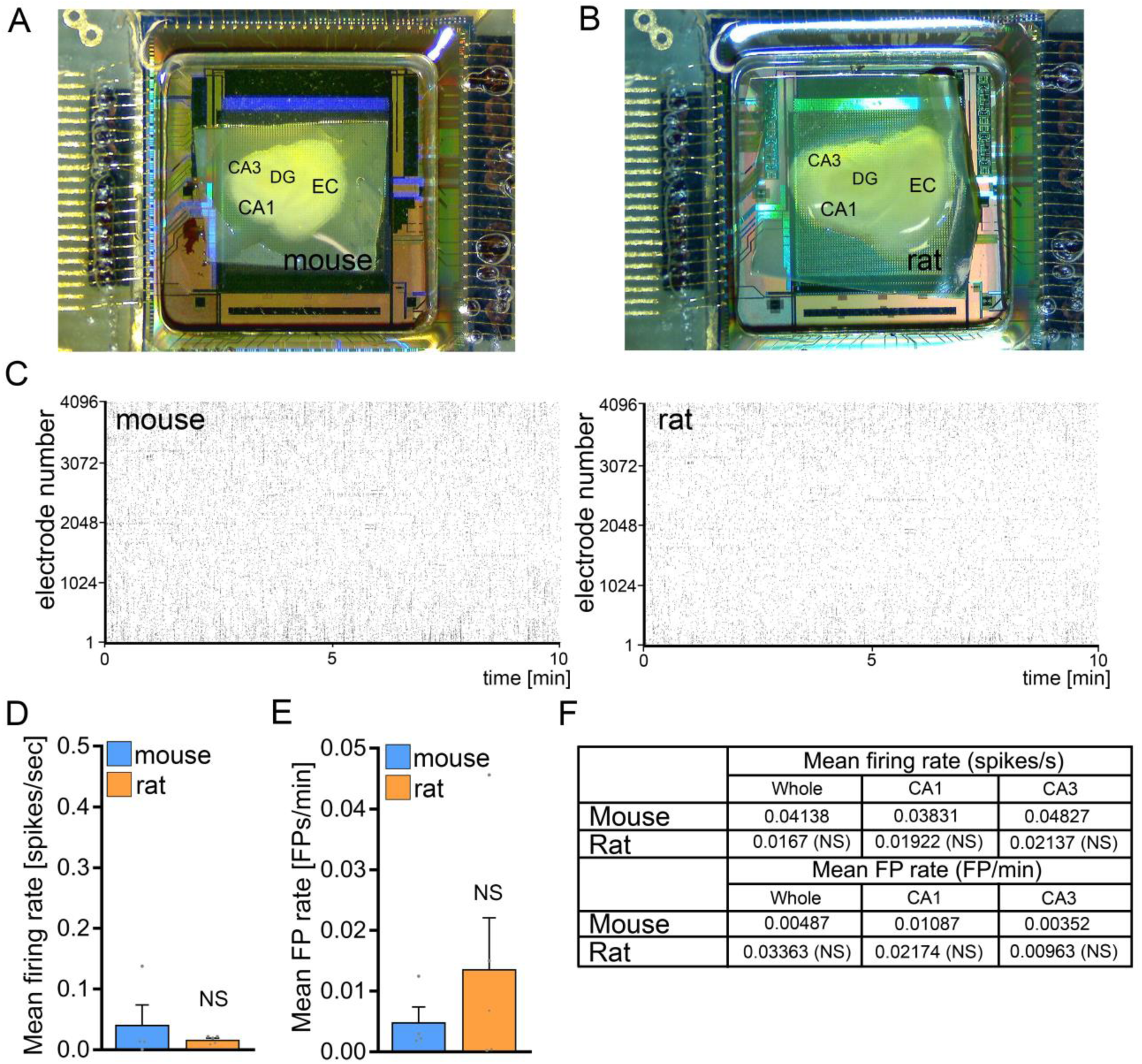
No significant differences in baseline network activity in mouse and rat tissue cultures. **(A-B)** Overview images of mouse and rat tissue culture on high-density multi electrode array chips. DG, Dentate gyrus; EC, entorhinal cortex; CA1 and CA3, *Cornu Ammonis* areas 1 and 3. **(C)** Raster plots of spikes during a 10-minute recording period in whole mouse and rat tissue cultures. **(D-F)** Group data of mean firing rate and mean field potential rate from mouse and rat tissue cultures (mouse, n = 4 cultures; rat, n = 5 cultures; Mann–Whitney test). Data are mean ± SEM. NS, Not significant.

### No significant differences in structural properties of cultured mouse and rat CA1 pyramidal neurons

To investigate whether differences in CA1 pyramidal neuron size and complexity could explain the variation in rMS outcome, we reconstructed biocytin-filled and streptavidin-A488 stained CA1 pyramidal neurons from both rat and mouse hippocampal tissue cultures and analyzed their dendrites and axons (Fig. 5). This was motivated by the observation that the brain sizes of mice and rats, as well as their tissue cultures, differ.

**Figure 5:**
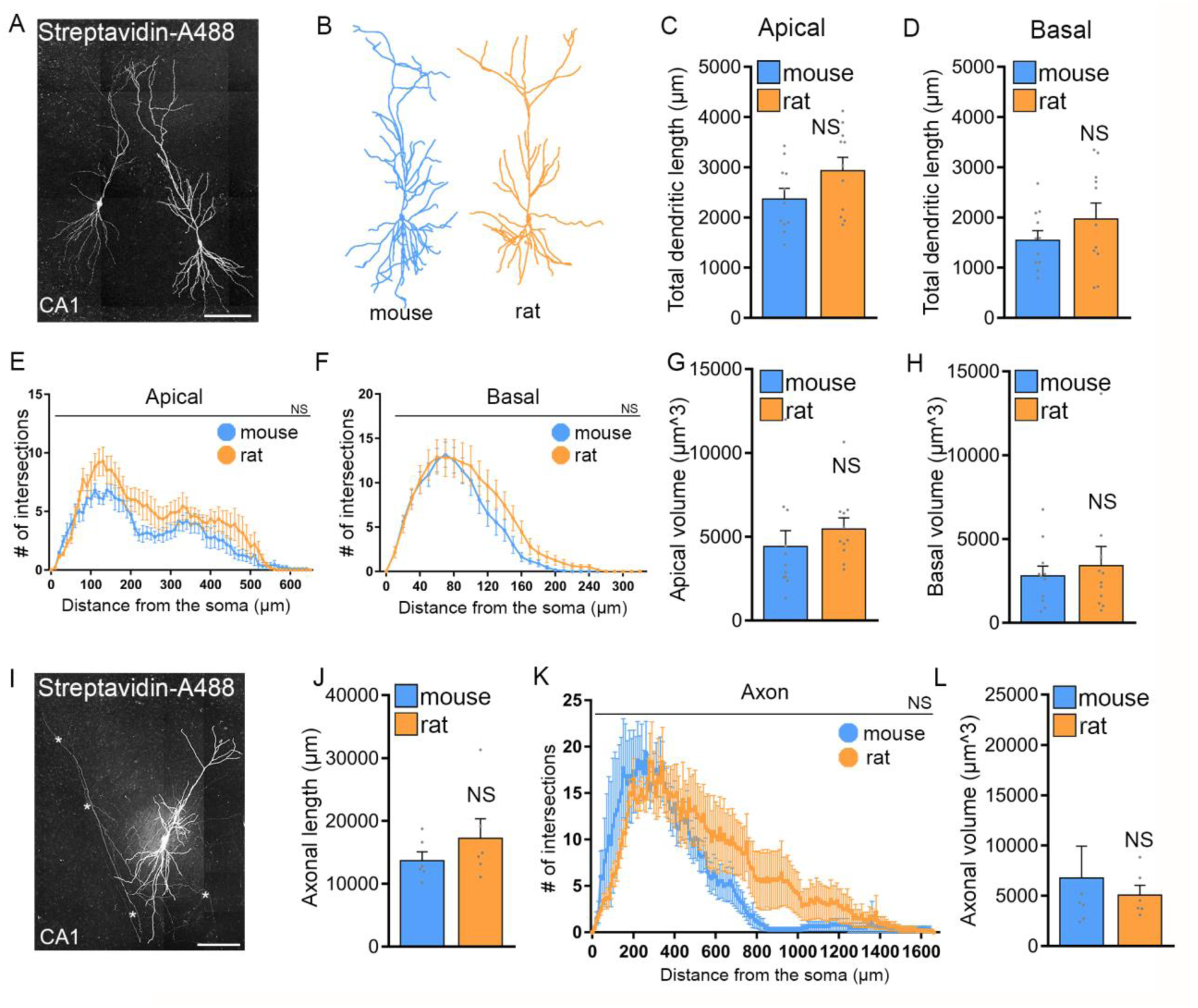
No significant morphological differences of CA1 pyramidal neurons in mouse and rat tissue cultures. **(A, B**) Examples of CA1 pyramidal neurons that were patched and filled with biocytin, later identified *post hoc* with streptavidin-A488, along with three-dimensional neuronal reconstructions of both mouse and rat CA1 pyramidal neurons. **(C-H)** Group data of mouse and rat apical and basal dendrites (mouse, n = 11 cells; rat, n = 11 cells; statistical comparisons for panels C, D, G and H were performed with Mann–Whitney test; statistical comparisons for panels E and F were performed with 2-way ANOVA). **(I)** Rat CA1 pyramidal neuron patched and filled with biocytin, identified *post hoc* with streptavidin-A488, and used for comprehensive neuronal reconstruction, encompassing dendritic and axonal neuronal structures. Scale bar, 50 μm. **(J-L)** Group data of mouse and rat axons (mouse, n = 6 cells; rat, n = 6 cells; statistical comparisons for panels J and L were performed with Mann–Whitney test; statistical comparisons for panel K were performed with 2-way ANOVA).

No significant differences were observed between the two groups in apical and basal dendritic length (Fig. 5C, D). Sholl and diameter/volume analyses (Fig. 5E-G) did not show any statistical significance between CA1 dendrites and their complexity of rat and mouse CA1 pyramidal neurons in entorhino-hippocampal tissue cultures. Similarly, no significant differences were observed when CA1 axons were reconstructed and compared in mouse and rat tissue cultures (Fig. 5I-L). We conclude, that structural properties of CA1 pyramidal neurons are not statistically different and cannot explain why the rat tissue cultures do not respond to 10 Hz rMS even when the E-field is closely matched based on E-field simulations.

### Realistic multiscale computer modeling predicts no major differences in rMS-induced depolarization of mouse and rat CA1 pyramidal neurons

We assessed the impact of rMS on CA1 pyramidal neurons through a multiscale computational model that connects the physical input parameters of rMS to dendritic and axonal morphologies (Fig. 6). This approach was necessary because our morphological analysis might not have encompassed distinctions pertinent to the neuronal activation induced by rMS.

**Figure 6:**
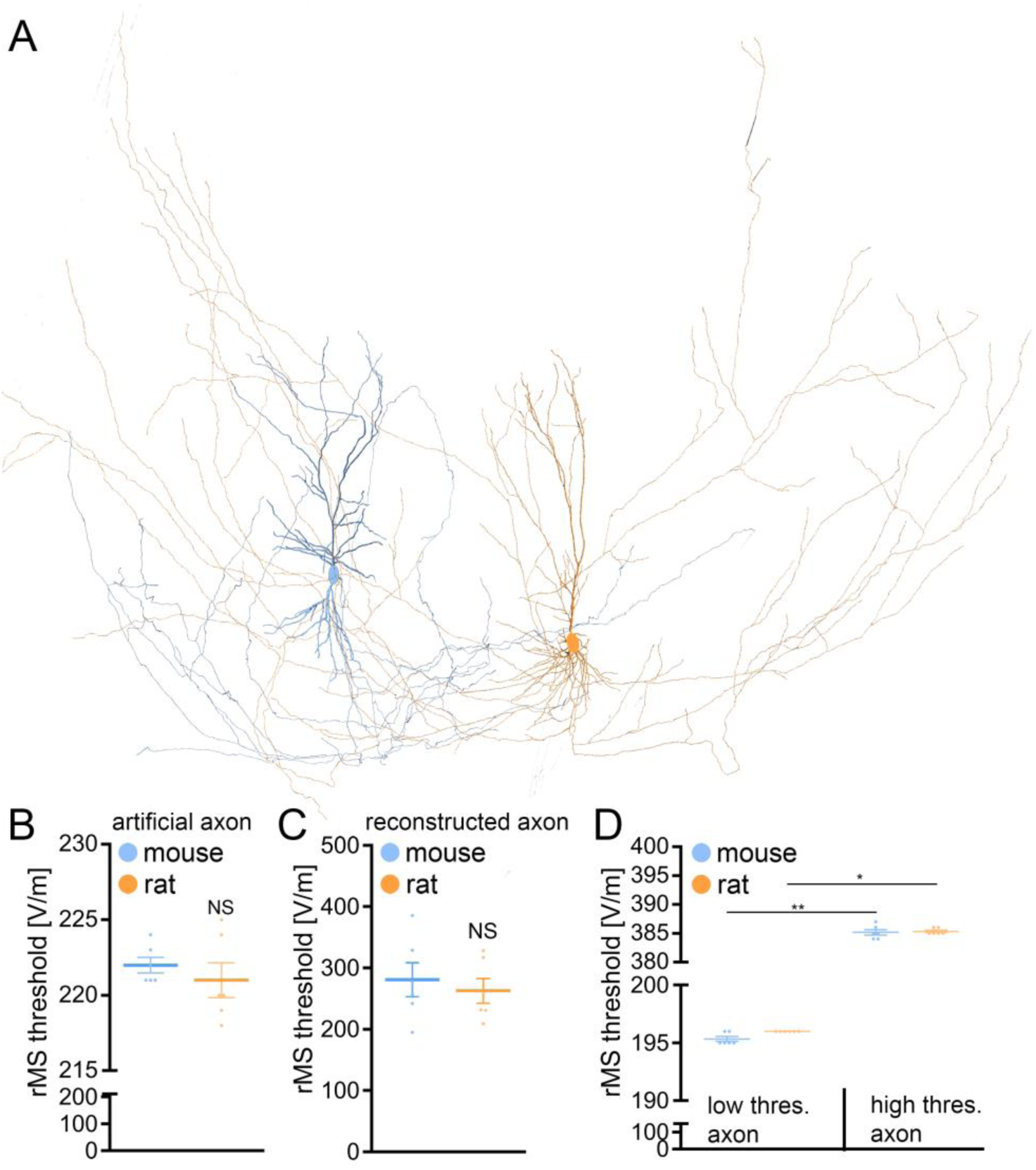
Multiscale computer modeling of electromagnetic stimulation. (A) Neuronal responses, i.e., changes in membrane voltage, to electromagnetic stimulation were modeled in realistic dendritic and axonal morphologies from reconstructed mouse and rat CA1 pyramidal neurons. (B) Group data of realistic dendritic morphologies with a standardized artificial axon (mouse, n = 6 cells; rat, n = 6 cells; Mann–Whitney test). (C) Group data of simulations with realistic dendritic and axonal morphologies (mouse, n = 6 cells; rat, n = 6 cells; Mann–Whitney test). (D) Group data for mouse and rat CA1 pyramidal neurons, categorizing those with axons exhibiting lowest (left) and highest (right) rMS depolarization thresholds (mouse, n = 6 cells; rat, n = 6 cells; Kruskal-Wallis test). Data are mean ± SEM. NS, Not significant.

When examining the dendritic architecture of CA1 neurons in mice and rats, and employing a standardized artificial axon across all cells (c.f., [16,29,30]), our simulations revealed no significant difference in the depolarization threshold elicited by rMS (Fig. 6A, B).

Subsequently, we investigated whether axonal morphologies might underlie the observed variability in our experimental outcomes. An additional series of simulations was conducted, this time integrating the authentic axonal morphologies of these neurons. Again, no significant differences in the depolarization thresholds were observed between the two groups (Fig. 6C).

A noteworthy insight emerged from these simulations: the axon’s influence is pivotal in establishing the rMS-induced depolarization threshold (Table 1). We followed up on this observation, by establishing connections between the axons responsible for the lowest and highest rMS depolarization thresholds across all mouse and rat cells. Indeed, a 2-fold difference in the depolarization thresholds was observed in these simulations across all reconstructed neurons (Fig. 6D). Yet, despite these simulations results, the dissimilarity in rMS-triggered plasticity between mouse and rat tissue cultures remained unresolved, eluding a complete explanation based solely on the interactions of dendritic and axonal morphologies.

**Table 1:** rMS-depolarization thresholds for individual cells with different axons attached.

|  | Axon 1 | Axon 2 | Axon 3 | Axon 4 | Axon 5 | Axon 6 | Axon 7 | Axon 8 | Axon 9 | Axon 10 | Axon 11 | Axon 12 |
| --- | --- | --- | --- | --- | --- | --- | --- | --- | --- | --- | --- | --- |
| Mouse cell 1 | 242 | 257 | 330 | 196 | 385 | 278 | 209 | 317 | 231 | 260 | 231 | 329 |
| Mouse cell 2 | 241 | 256 | 329 | 195 | 384 | 278 | 209 | 316 | 231 | 260 | 231 | 328 |
| Mouse cell 3 | 241 | 256 | 329 | 195 | 384 | 278 | 209 | 316 | 231 | 260 | 231 | 328 |
| Mouse cell 4 | 242 | 258 | 330 | 195 | 385 | 278 | 209 | 317 | 231 | 260 | 232 | 328 |
| Mouse cell 5 | 241 | 255 | 330 | 195 | 386 | 278 | 209 | 317 | 231 | 260 | 231 | 328 |
| Mouse cell 6 | 241 | 257 | 330 | 196 | 387 | 278 | 209 | 317 | 231 | 260 | 232 | 329 |
| Rat cell 1 | 242 | 257 | 330 | 196 | 385 | 278 | 209 | 317 | 231 | 260 | 231 | 329 |
| Rat cell 2 | 242 | 258 | 330 | 196 | 386 | 278 | 209 | 317 | 231 | 260 | 231 | 329 |
| Rat cell 3 | 243 | 258 | 330 | 196 | 385 | 278 | 209 | 317 | 231 | 261 | 232 | 329 |
| Rat cell 4 | 242 | 257 | 330 | 196 | 385 | 278 | 209 | 317 | 231 | 260 | 231 | 329 |
| Rat cell 5 | 242 | 258 | 331 | 196 | 386 | 278 | 209 | 317 | 231 | 260 | 232 | 329 |
| Rat cell 6 | 241 | 258 | 330 | 196 | 385 | 278 | 209 | 317 | 231 | 260 | 231 | 328 |

Values represent rMS-depolarization threshold in V/m

### Active and passive membrane properties reveal differences in excitability between mouse and rat CA1 pyramidal neurons

Next, active and passive membrane properties were recorded from CA1 pyramidal neurons and analyzed. Indeed, this set of experiments identified significant differences in the passive and active properties between mouse and rat CA1 pyramidal neurons (Fig. 7).

**Figure 7:**
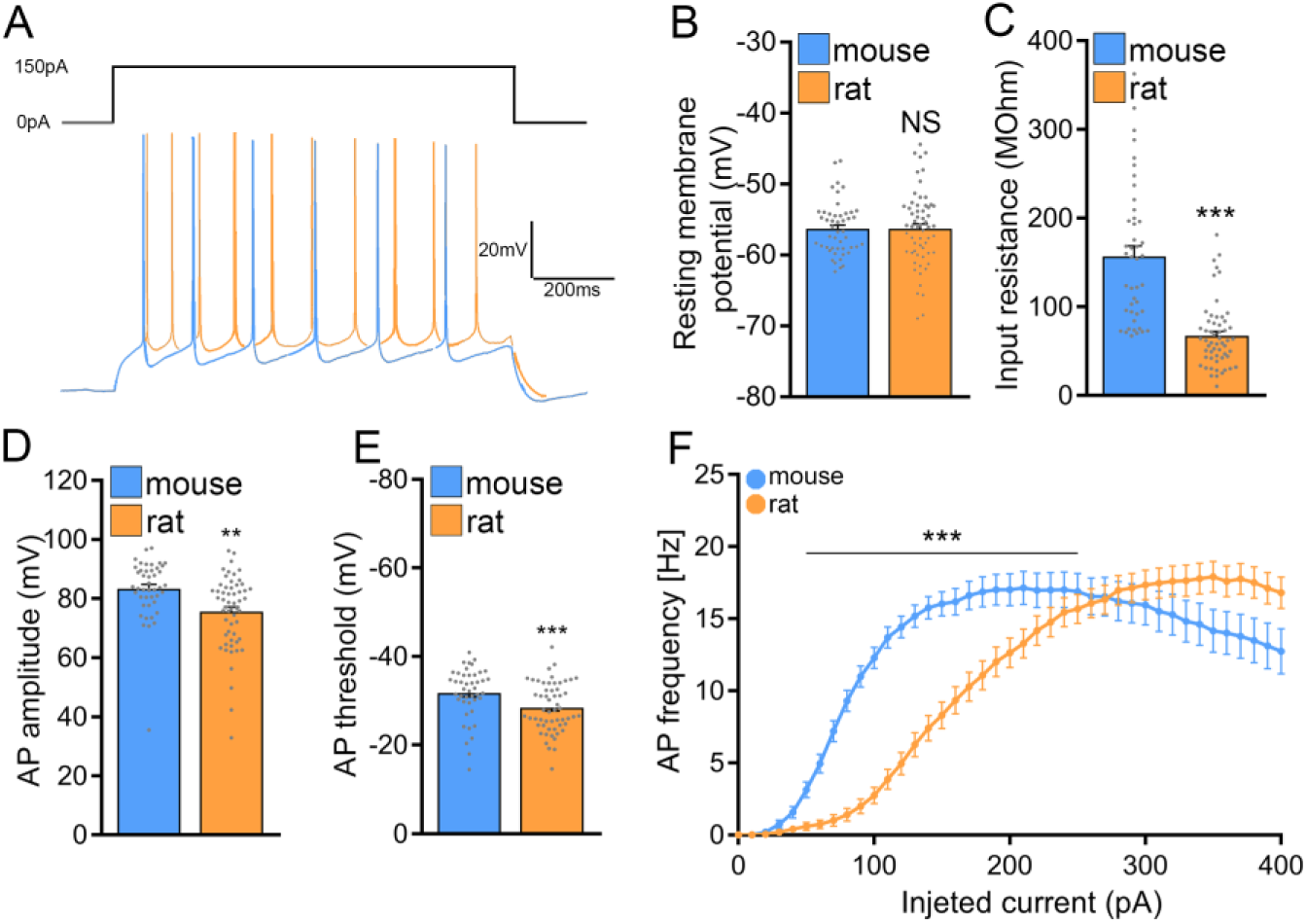
Rat CA1 pyramidal neurons exhibit lower excitability in comparison to mice. **(A**) Sample traces from input-output recordings of CA1 pyramidal neurons of mouse and rat tissue cultures. **(B, C)** Group data of resting membrane potential and input resistance from mouse and rat CA1 pyramidal neurons (mouse, n = 44 cells; rat, n = 56 cells; Mann–Whitney test). **(D, E)** Group data of action potential (AP) amplitude and threshold from mouse and rat CA1 pyramidal neurons (mouse, n = 44 cells; rat, n = 56 cells; Mann–Whitney test). **(F)** I-F curve of CA1 pyramidal neurons of mouse and rat tissue cultures (mouse, n = 52 cells; rat, n = 63 cells; 2-way ANOVA). Data are mean ± SEM. NS, Not significant. **p < 0.01. ***p < 0.001.

The input resistance of mouse CA1 pyramidal neurons was significantly higher as compared to rat CA1 pyramidal neurons (mouse: 156.8 ± 11.65 MOhm and rat: 67.25 ± 4.909 MOhm; Mann-Whitney test; p < 0.001; U = 279), while the cells of both mice and rats were resting at comparable membrane potentials (Fig. 7A-C). Consistently, the current-voltage (I/V) curves demonstrated that depolarizing mouse CA1 pyramidal neurons is more straightforward compared to cultured rat CA1 pyramidal neurons.

Looking at the active membrane properties (Fig. 7D-F) a similar trend was observed with the most striking differences being in the action potential induction threshold (mouse: -31.81 ± 0.877 mV; rat: -28.47 ± 0.744 mV; Mann-Whitney test; p = 0.0021; U = 794) and the first spike latency (mouse: 419.8 ± 56.03 ms; rat: 715 ± 77.36 ms; Mann-Whitney test; p = 0.0074; U = 15; data not shown). Figure 7F, shows that current injections produced stronger responses in mouse CA1 pyramidal neurons than in rat neurons, i.e., higher action potential frequencies at a lower current injection. These results indicated that mouse CA1 pyramidal neurons are more excitable than rat neurons, suggesting that higher stimulation intensities may be needed to induce rMS-induced plasticity in rat tissue cultures.

### 60 % MSO induces rMS-mediated plasticity in rat organotypic tissue cultures

Subsequently, we tested whether a 10 Hz stimulation protocol applied at a higher intensity would induce plasticity in rat CA1 pyramidal neurons. Indeed, when rat tissue cultures were stimulated with 10 Hz rMS at 60 % MSO a robust increase in the mean mEPSC amplitude was detected (Fig. 8A), similar to what we observe in the mouse cultures stimulated at 50 % MSO (cf. Fig. 1E and Fig 3D). In addition, a significant reduction in mean mIPSC amplitude was evident 2 – 4 h after rMS stimulation at 60 % MSO in a different set of rat tissue cultures (Fig. 8B; c.f., Fig. 1G). These results demonstrate that rat CA1 pyramidal neurons do express rMS-induced plasticity, but require a higher stimulation intensity for rMS-induced potentiation of excitatory synapses and depression of inhibition to occur.

**Figure 8:**
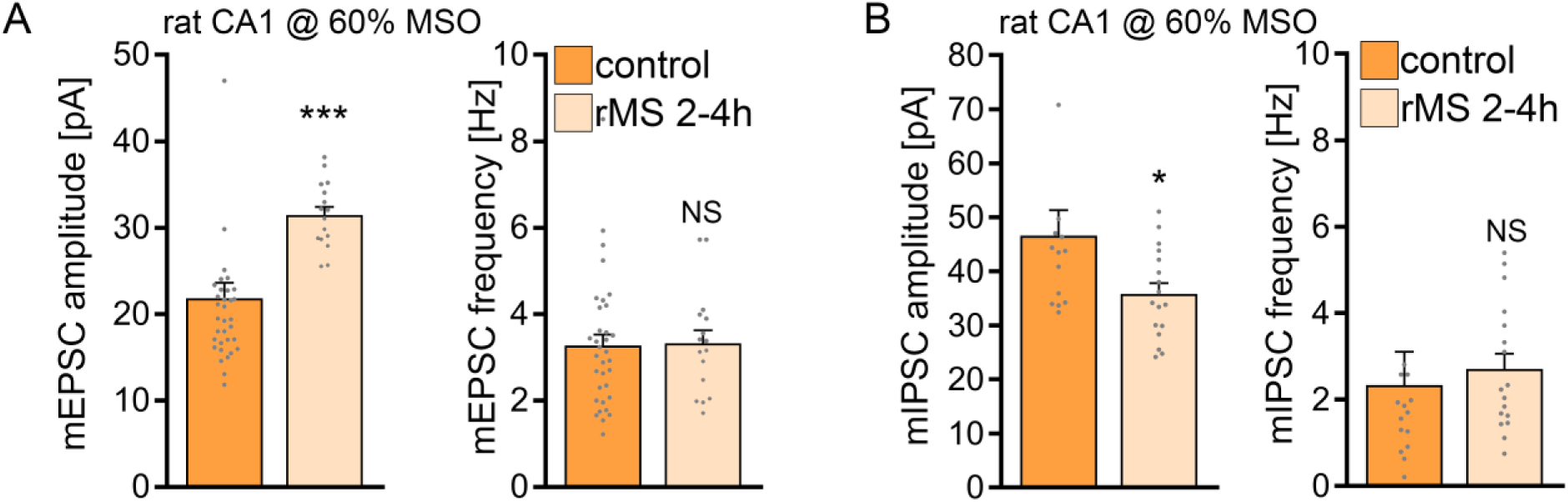
10 Hz repetitive magnetic stimulation (rMS) at 60 % MSO induces synaptic plasticity in rat CA1 pyramidal neurons. (A) Group data of AMPA receptor mediated mEPSCs recorded from rat CA1 pyramidal neurons from sham-(control) and rMS-stimulated cultures (control, n = 34 cells; rMS, n = 16 cells; Mann– Whitney test). (C, D) Sample traces and group data of miniature inhibitory post synaptic currents (mIPSCs) recorded from rat CA1 pyramidal neurons from sham-(control) and rMS-stimulated cultures (control, n = 14 cells; rMS, n = 17 cells; Mann–Whitney test. One data point outside of axis limits in mIPSC amplitude and frequency respectively). Data are mean ± SEM. NS, Not significant. *p < 0.05. ***p < 0.001.

## DISCUSSION

In this study, we explored the factors influencing the threshold for 10 Hz rTMS-induced synaptic plasticity. Using mouse and rat entorhino-hippocampal slice cultures, we investigated neuronal structure, excitability, and network activity. In mouse CA1 pyramidal neurons, we confirmed the well-known potentiation of excitatory synapses and depression of inhibitory synapses, highlighting robust rTMS-induced synaptic plasticity under controlled conditions. However, despite similar neuronal morphology and network activity in rat CA1 pyramidal neurons, standardizing electric fields through prospective modeling did not produce the same biological effect. Adjusting the stimulation protocol to account for rat neurons’ lower excitability led to comparable synaptic changes. These results emphasize that electric field standardization alone cannot predict rTMS effects, necessitating realistic compartmental models of cellular properties in different brain regions for accurate predictions.

Over the past decade, the utilization of rTMS has experienced a significant surge in both research and clinical domains [3,37–41]. Consequently, extensive efforts have been dedicated to identify the crucial parameters that influence the effects of rTMS on brain tissue [25,40,42,43]. Among these parameters, the induced electric field emerges as a pivotal factor directly shaping the impact of rTMS on cortical tissue [44]. While advancements in computational tools have enabled the calculation of rTMS-induced electric field [45], these models have primarily relied on mesoscopic structural parameters of the targeted stimulation area, i.e., head and brain geometries. In recent years, there has been a growing adoption of multi-scale modeling approaches to investigate the impact of TMS on individual neurons [29–31,46]. Notably, these neuronal models are being integrated into mesoscopic brain models, enabling exploration of the effects of cortical folding and the precise positioning of neurons, such as distinguishing between the gyral crown and gyral groove, in individual subjects [31,47–49]. While these models represent a significant advancement toward standardization and precision medicine in the field, it is increasingly evident that solely modeling electric fields and their interactions with individual neuronal morphologies (derived from animal models) may not be sufficient to standardize the biological effects of rTMS across various brain regions and individuals [49]. The findings from this cross-species study present experimental evidence, underscoring the insufficiency of meticulous experimental standardization and electric field modeling in guaranteeing robust biological effects of rTMS. Notably, computational modeling showed weaker induced electric fields in rat tissue cultures despite their size difference compared to mouse tissue cultures. Even when efforts were made to match electric fields, the plasticity effects in rat cultures could not be reproduced.

In this context, it is crucial to highlight that our experiments revealed no statistically significant morphological differences between the cultured CA1 pyramidal neurons of mice and rats. The comprehensive analysis of both apical and basal dendrites demonstrated comparable total dendritic length, complexity, and overall volume in both rat and mouse pyramidal neurons of organotypic tissue cultures. These results align with previously published data that compared mouse and rat hippocampal CA1 neurons in acute slice preparations [50]. However, it is worth noting that the total volume of these cells, apart from the observed morphologies features, was found to be higher in rat slices. Though differences between acute brain slices and tissue cultures could contribute to the observed discrepancy, it is crucial to highlight the key advantage of tissue cultures. Using 3-week-old tissue cultures enabled us to investigate neurons within brain tissue that had not undergone acute slicing immediately before experimental assessment. This allowed us to study undamaged CA1 pyramidal neurons and enabled us to generate detailed morphological reconstructions, encompassing both dendrites and axons. Specifically, complete reconstructions of axons are of utmost importance for precise evaluation of rTMS outcomes, considering their substantial interaction with the electric field [51]. Previous studies, including our own work, often relied on artificial or simplified axon morphologies [16, 29–31]. Importantly, our investigation revealed no significant differences in axons of cultured CA1 neurons between mice and rats. This finding suggests that the observed inability of rat CA1 neurons to exhibit synaptic plasticity cannot be trivially attributed to differences in axon morphology.

Nevertheless, our simulations identified axons that are twice as effective at depolarizing neurons, irrespective of soma and dendrite shapes. This emphasizes the need for a systematic assessment of various axonal morphologies in rTMS-induced synaptic plasticity, also considering factors like myelination and the role of oligodendrocytes. We propose the possibility of “super-responder cells” within complex cortical networks–cells highly responsive to rTMS at specific stimulation intensities. This notion finds support in the observation that not all neurons of this and our previous studies (c.f., [11,13,16,36]) displayed elevated mEPSC amplitudes or decreased mIPSC within the 2 – 4 h following stimulation.

The results of the present study suggest that understanding the differing effects of rTMS on mouse and rat CA1 pyramidal neurons requires considering their intrinsic cellular properties. Consistent with prior research on rat and mouse slices [50], our study shows that rat CA1 pyramidal neurons have a higher action potential threshold compared to mice, making them less excitable. Notably, we found that rat CA1 neurons have lower input resistance than mouse neurons, further highlighting reduced excitability in rat neurons. However, it’s worth noting that a study by Routh and colleagues in 2009 reported similar input resistance between the two species [50], potentially due to differences in acute slices prepared from adult animals and organotypic tissue cultures.

Do morphological and biophysical properties alone predict rTMS outcomes adequately? Additional factors, like neuromodulators such as dopamine, serotonin, and noradrenaline, influence cortical excitability, impacting how neurons respond to rTMS and altering plasticity threshold, magnitude, and direction. [52–57]. Furthermore, neuromodulators can impact the capacity of neurons to express plasticity without affecting excitability and other baseline functional and structural properties, a phenomenon known as metaplasticity [58,59]. It is important to also note that non-neuronal cells can significantly influence the capacity of neurons to express synaptic plasticity [60–65]. Our prior work has provided evidence that cytokines derived from microglia play a crucial role in facilitating rTMS-induced plasticity [16]. Finally, the impact of network activity on plasticity thresholds and the outcome of rTMS must be considered. These factors collectively underscore the multifaceted nature of the processes involved in influencing and modulating the outcomes of rTMS-induced plasticity. Organotypic slice cultures serve as valuable tools for investigating these and other aspects of rTMS-induced plasticity, highlighting the necessity for rigorously validated computer models that link the induced electric fields with biophysically realistic neurons and networks. These models hold the potential to predict the biological outcomes of rTMS, offering valuable insights into its effects and guiding the adaptation of stimulation protocols to achieve consistent desired effects across different brain regions and individuals.

## AUTHOR CONTRIBUTIONS

**Christos Galanis:** Conceptualization; Methodology; Investigation; Formal analysis; Data Curation; Visualization; Supervision; Writing – Original Draft, Review & Editing. **Lena Neuhaus:** Investigation; Formal analysis. **Nicholas Hananeia:** Software; Investigation; Formal analysis; Data Curation. **Zsolt Turi**: Software; Investigation; Formal analysis; Data Curation. **Peter Jedlicka:** Methodology; Resources; Software; Data Curation; Supervision. **Andreas Vlachos:** Conceptualization; Methodology; Data Curation; Supervision; Writing – Original Draft, Review & Editing; Project administration; Funding acquisition.

## AKNOWLEDGMENTS

**Acknowledgements:** We thank Susanna Glaser for skillful assistance in tissue culturing.

